# A Newly Identified Role for IL-22 in Bladder Immunity

**DOI:** 10.1101/2020.08.17.255273

**Authors:** Jared Honeycutt, Loc Le, Yi-Ju Hsieh, Michael H. Hsieh

**Affiliations:** Stanford Immunology, Stanford University, Stanford, CA; Biomedical Research Institute, Rockville, MD; Children’s National Health System, Washington, DC; The George Washington University, Washington, DC

## Abstract

Bacterial and parasitic urinary tract infections(UTI) affect many, but our understanding of associated immunity remains poor. Prior work indicates interleukin-22(IL-22) is a key cytokine during infection of other epithelia. We hypothesized IL-22 is crucial during UTI.

IL-22-null(KO) and wild-type(wt) mice underwent bladder wall injection with *S. haematobium* eggs or transurethral infection with uropathogenic *E. coli* UTI89.

IL-22 and its soluble binding protein, IL-22BP, were increased in the bladder after *S. haematobium* egg injection. Genes typically induced by IL-22(CXCL2, REG3G[antimicrobial defense protein]], S100A8, S100A9, and Areg) were expressed at higher levels following *S. haematobium* egg injection. IL-22 stimulation of bladder tissue induced REG3B expression. IL-22 receptor alpha-1 expression was detectable in the urothelium by immunofluorescence and qPCR.

Injection of *S. haematobium* eggs into IL-22-KO vs wt mice triggered differential expression of genes related to transferase activity(transferring alkyl or aryl groups) and epithelial cell development. Numerous uroplakin genes were downregulated in egg-injected, IL-22-KO mice versus their wt counterparts. These decreases in uroplakin expression suggest that, as in the gut, IL-22 is important for replenishing the epithelium during infection-related injury.

Following transurethral infection with UTI89, IL-22-KO mice had lower bacterial counts in their urine, bladder, and kidneys. Giving stabilized IL-22 cytokine (IL-22-Fc) to UTI89-infected mice led to higher kidney bacterial counts and increased morbidity.

Our data suggest that IL-22 is indeed important in urinary tract immunity, and may interfere with clearance of bacteria from the urinary tract, potentially through its role in maintenance of mature urothelium.

## Introduction

In contrast to what is known of the gut and lung mucosa, it is unclear how inflammatory pathways in the urinary tract contribute to host protection and pathology during the parasitic and bacterial infections that occur at this site. This is a key question, given that: 1)) bacterial UTI affect half of all girls and women at least once during their lifetimes^1^; 2) urogenital schistosomiasis, a parasitic UTI caused by *Schistosoma haematobium* worms, affects 112 million people globally^2^; and 3) urogenital schistosomiasis induces major changes in the bladder urothelium, including hyperplasia, ulceration with egg expulsion and hematuria, squamous metaplasia, and frank carcinogenesis (reviewed by Honeycutt et al. ^3^).

Recent reports indicate that interleukin-22 (IL-22) is a key cytokine supporting epithelial immunity, while its improper regulation may contribute to pathologies such as fibrosis and cancer. We hypothesized that IL-22 may play an important role in bladder immunity and homeostasis during *Schistosoma haematobium* and *E.coli* infections. Moreover, we postulated that IL-22 may have different effects on the bladder versus other endodermal organs, given that the bladder: 1) is exposed to a spectrum of microbiota that only partially overlaps with that found in the gut and lung; and 2) has no major submucosal lymphoid structures, except during chronic inflammation.

IL-22 is produced by activated dendritic cells (DC), innate lymphoid cells (ILC), lymphoid tissue inducer (LTi) cells, natural killer (NK), NK T cells, and T cells, and initiates innate immune responses against a range of pathogens (reviewed by Rutz et al.^4^). Responsiveness to IL-22 is limited to parenchymal cells such as hepatocytes, keratinocytes, and the epithelial cells lining endodermal derivatives (i.e., the gut and respiratory tract), because only these cells express the appropriate receptor complex (heterodimers of IL-[IL--10R2, IL-22R). Upon binding this receptor complex, IL-22 promotes epithelial regeneration and secretion of antimicrobial peptides (reviewed by Eidenschenk et al.^5^).

Herein we utilized experimental mouse models of bacterial UTI and urogenital schistosomiasis to dissect the roles of IL-22 in bladder inflammation induced by a range of uropathogens. We present evidence suggesting that IL-22 indeed play a key role in urinary immunity to both prokaryotic and eukaryotic pathogens.

## Results

We dissected the urothelial lining from the bladder wall of naïve mice and used qPCR to measure expression of IL22Rα1 and other genes associated with mature urothelium (*Upk3a, Upk1b)* relative to the muscular bladder wall (Figure 1A). We also evaluated protein expression in the urothelium by immunofluorescence microscopy of frozen sections of naïve bladder tissue co-stained with monoclonal antibodies to IL22Rα1 and the urothelial marker cytokeratin-5 (Figure 1B).

**Figure 1.**
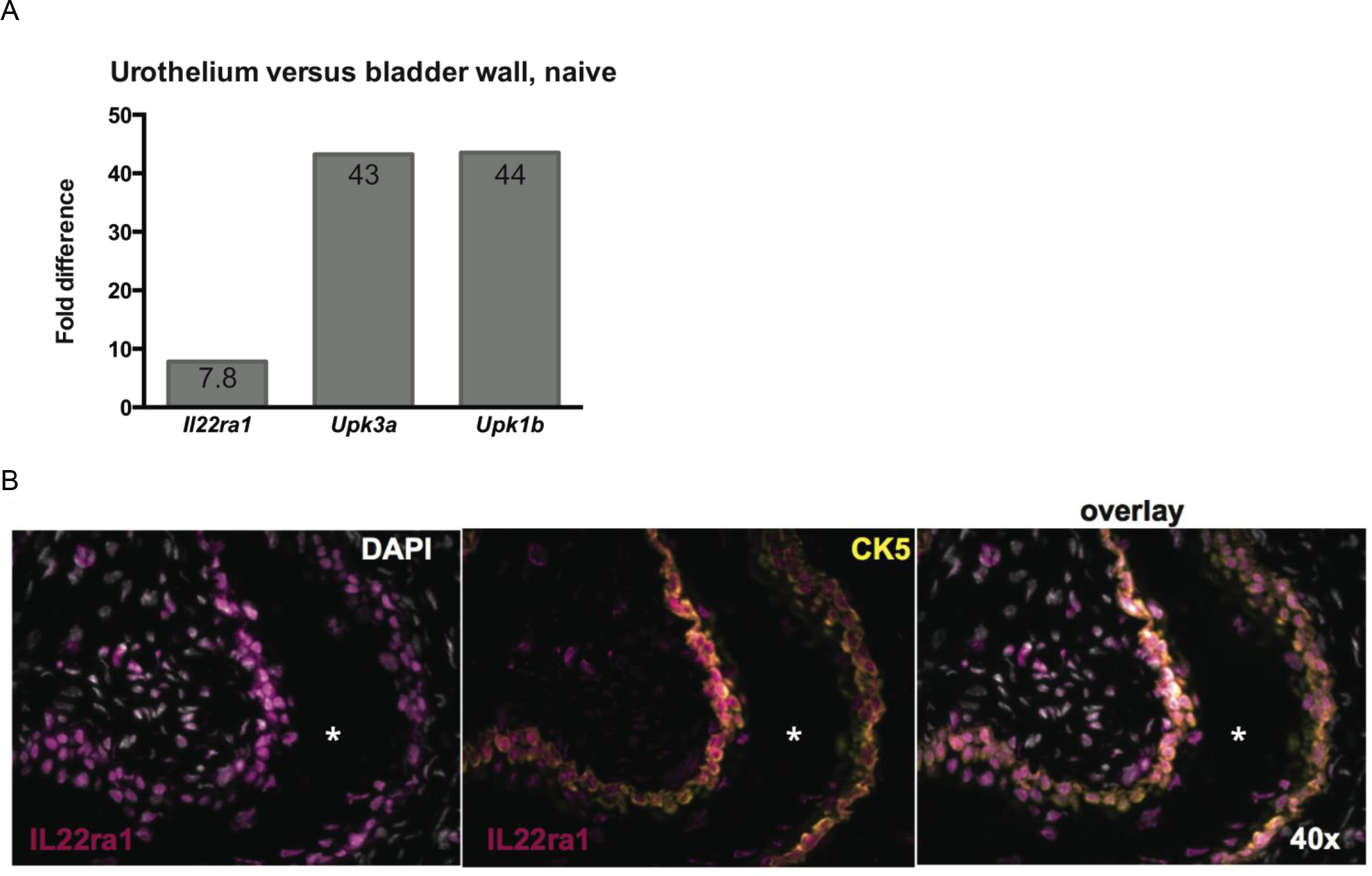
Expression of IL22 receptor alpha-1 in the mouse bladder. (A) The urothelium was manually microdissected from the muscular bladder wall of naïve C57BL/6J female mice. Relative expression of *IL22ra1* and the urothelial-specific genes *Upk3a* and *Upk1b* were measured by qPCR with *Gapdh* as the reference gene. (B) Naïve C57BL/6J female mouse bladders were frozen and sectioned, staining for IL22Rα1, cytokeratin 5 (Ck5 - a marker for mature urothelium), and DAPI to detect nuclei. * indicates lumen of bladder.

Using a model of chronic bladder inflammation induced by *Schistosoma haematobium* egg injection into the bladder wall, we observed increased transcription of *Il22* and its soluble regulator *Il22bp* in the bladder tissues of egg-injected mice. This occurred within the context of the previously observed type 2 immune response induced by egg injection (Figure 2A). Further qPCR measurements revealed that several genes classically induced by IL-22 were also upregulated in egg-injected BALB/c mice (Figure 2B).

**Figure 2.**
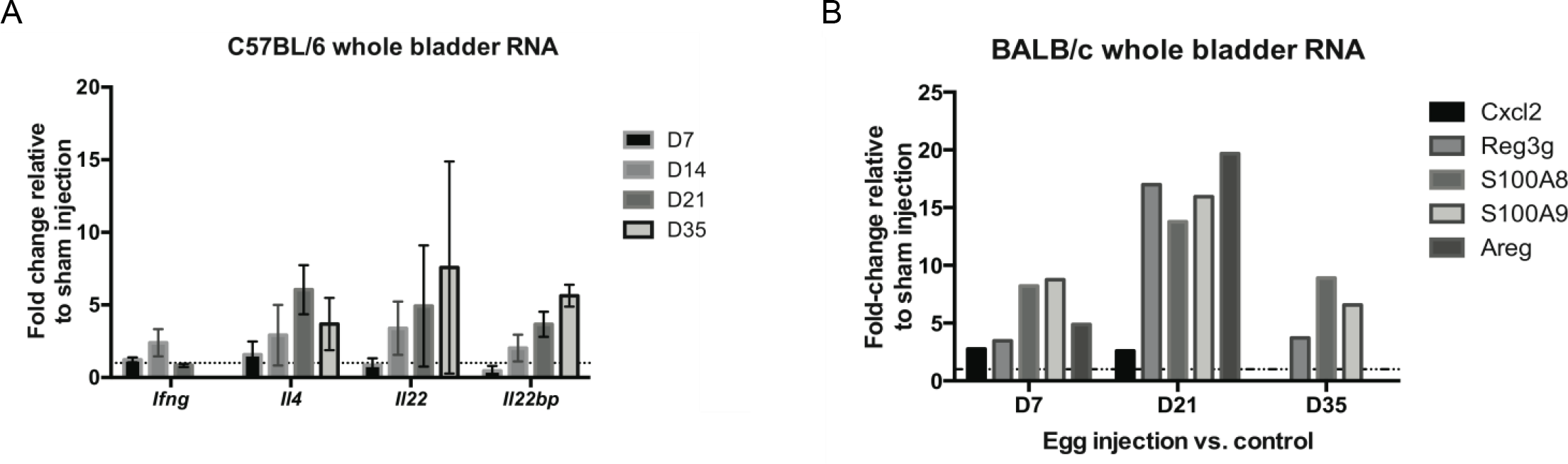
IL-22 and target gene expression in *S. haematobium* egg injected bladder. (A) C57BL/6J female mouse bladders were injected with *S* .*haematobium* eggs or sham injections and transcript levels were compared by qPCR at the indicated timepoints (n=4 per group at each timepoint). (B) Expression of anti-microbial genes that are inducible by IL-22 in BALB/c female egg-injected mice relative to sham injected mice at indicated days post egg injection.

We sought to evaluate the effects of IL-22 during chronic bladder inflammation, given its previously documented role in fibrosis and cancer in the liver and gut, respectively^6,7^. We induced chronic bladder inflammation by injecting *Schistosoma haematobium* eggs into the bladder walls of *IL22*-/-and wild type (WT) littermate mice (n=4 per group, 2 males and 2 females per group). At day 21 post egg injection, whole bladder RNA was prepared and samples were compared by gene expression microarrays. Analysis of expression differences identified significant changes in two gene ontology (GO) classes: epithelial development and transferase activity. Genes related to urothelial maturation (e.g., uroplakins) and the Wnt/Hedgehog pathways were downregulated in egg-injected, *IL22*-/-mice relative to their WT littermates (Figure 3A). The alterations in several Wnt/Hedgehog pathway genes were validated by qPCR (Figure 3B). There were also significant reductions in expression of multiple glutathione transferase genes in *IL22*-/-egg-injected bladders compared to WT (Figure 3C). In line with the significant expression changes observed, representative histology suggested that the urothelium overlying bladder granulomas exhibited more ulceration in *IL22*-/-compared to WT mice (Figure 4).

**Figure 3.**
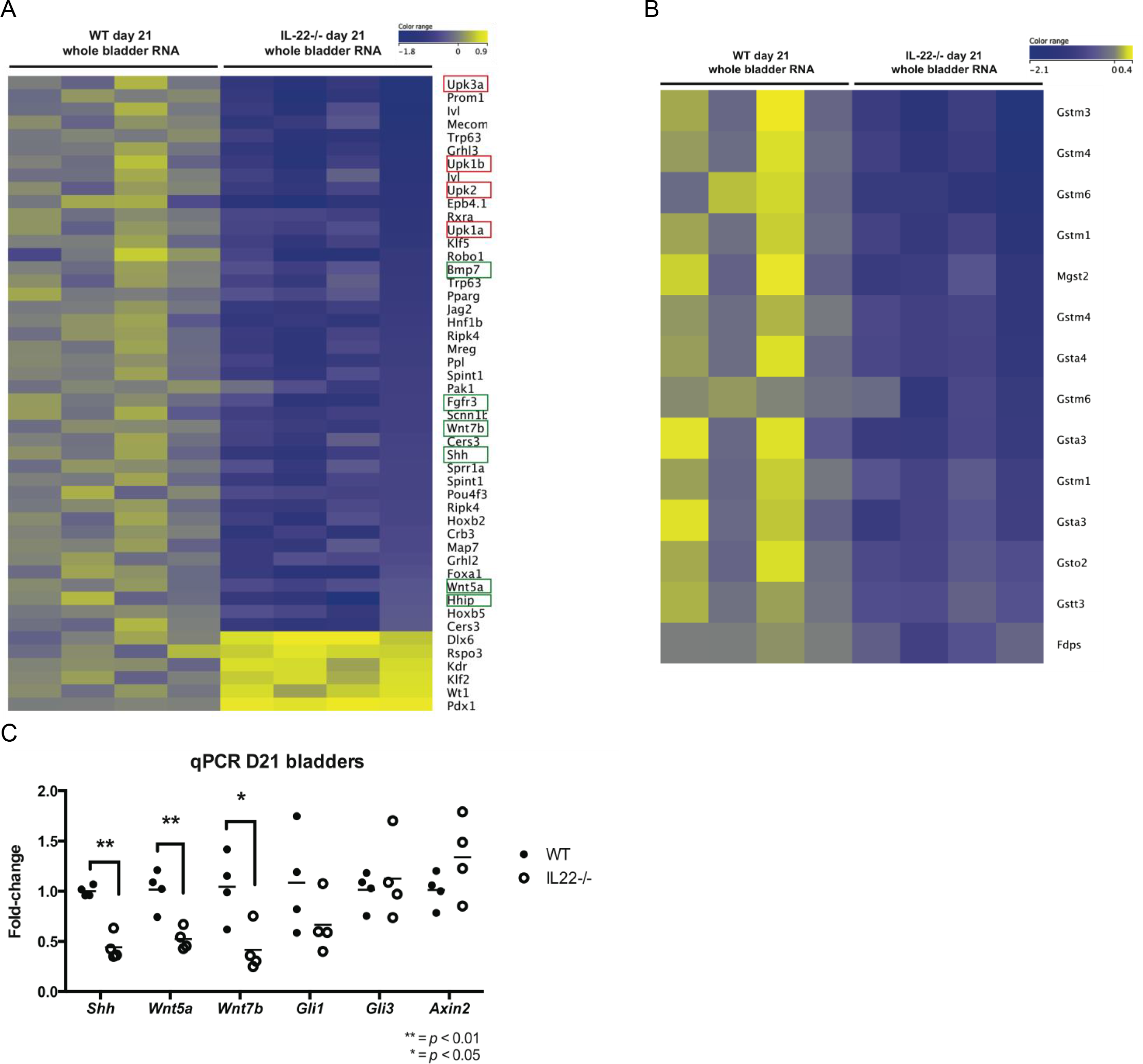
Comparison of *S. haematobium* induced gene expression in bladders of WT and *IL22-/-* mice. (A) Microarray was performed on whole bladder tissue RNA 21 days after egg injection. Expression differences related to gene ontology (GO) term “epithelial development”. Urothelial related genes outlined in red. Regenerative pathway genes outlined in green. (B) Expression differences related to GO term “transferase activity”. (C) qPCR validation of select Shh and Wnt pathway genes with *Hprt* as the reference gene.

**Figure 4.**
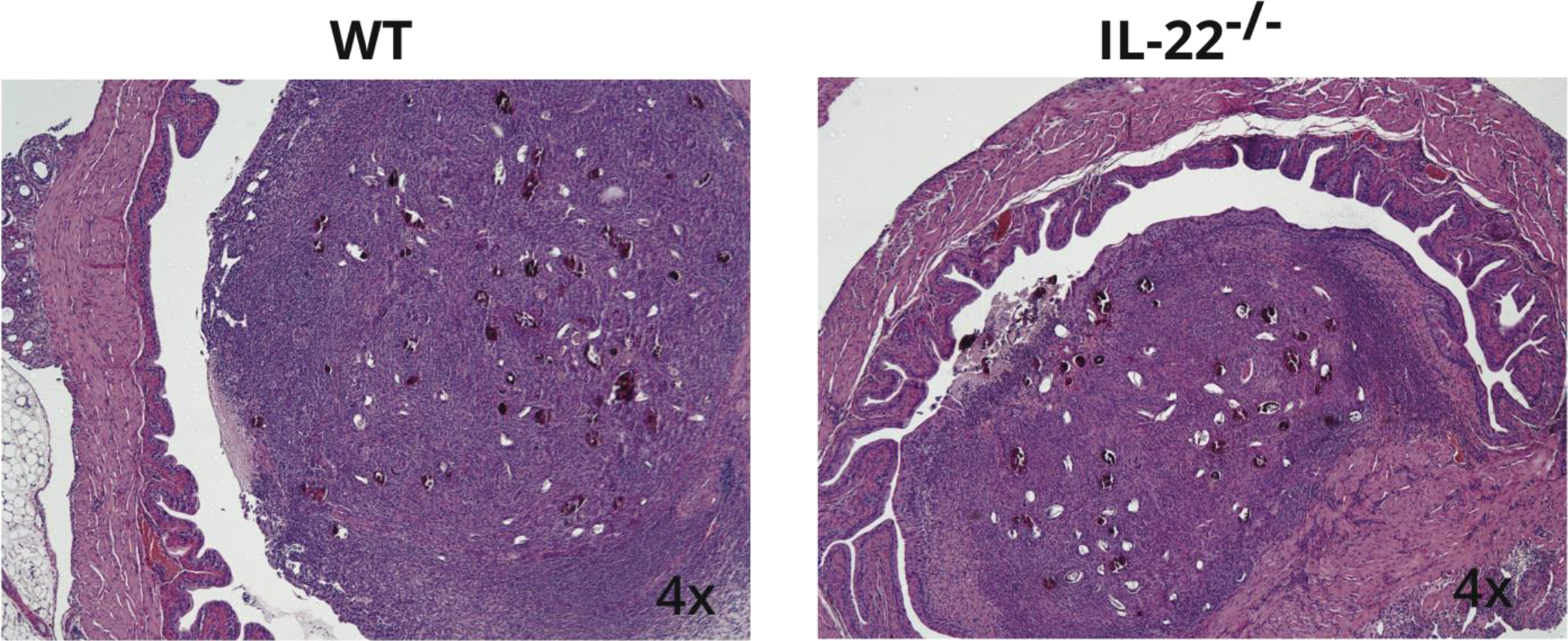
Representative histology from *IL22-/-* and WT littermates. *IL22-/-* (n=2) and WT (n=4) mice received a bladder wall injection of 3000 *Schistosoma haematobium* eggs and at day 21 bladders were fixed in formalin and embedded in paraffin for H&E staining. A number of noticeably large urothelial lesions (areas of ulceration) were noted in the IL-22-/-bladders.

IL22 plays an important role in successful responses to bacterial infections at several mucosal sites. We tested the role of IL22 in acute bacterial urinary tract infection using the uropathogenic *E. coli* strain, UTI89. Two days after infection, *IL22-/-* mice showed no statistically different response to UTI89, though the trend was lower in this high variance model, warranting further replication (Figure 5A). We also tested whether administration of exogenous IL-22 prior to and during infection had an effect on bacterial burdens (Figure 5B). In C57BL/6 female mice, there was a significant difference in kidney CFUs at day 2 including increased morbidity in IL22-Fc treated mice.

**Figure 5.**
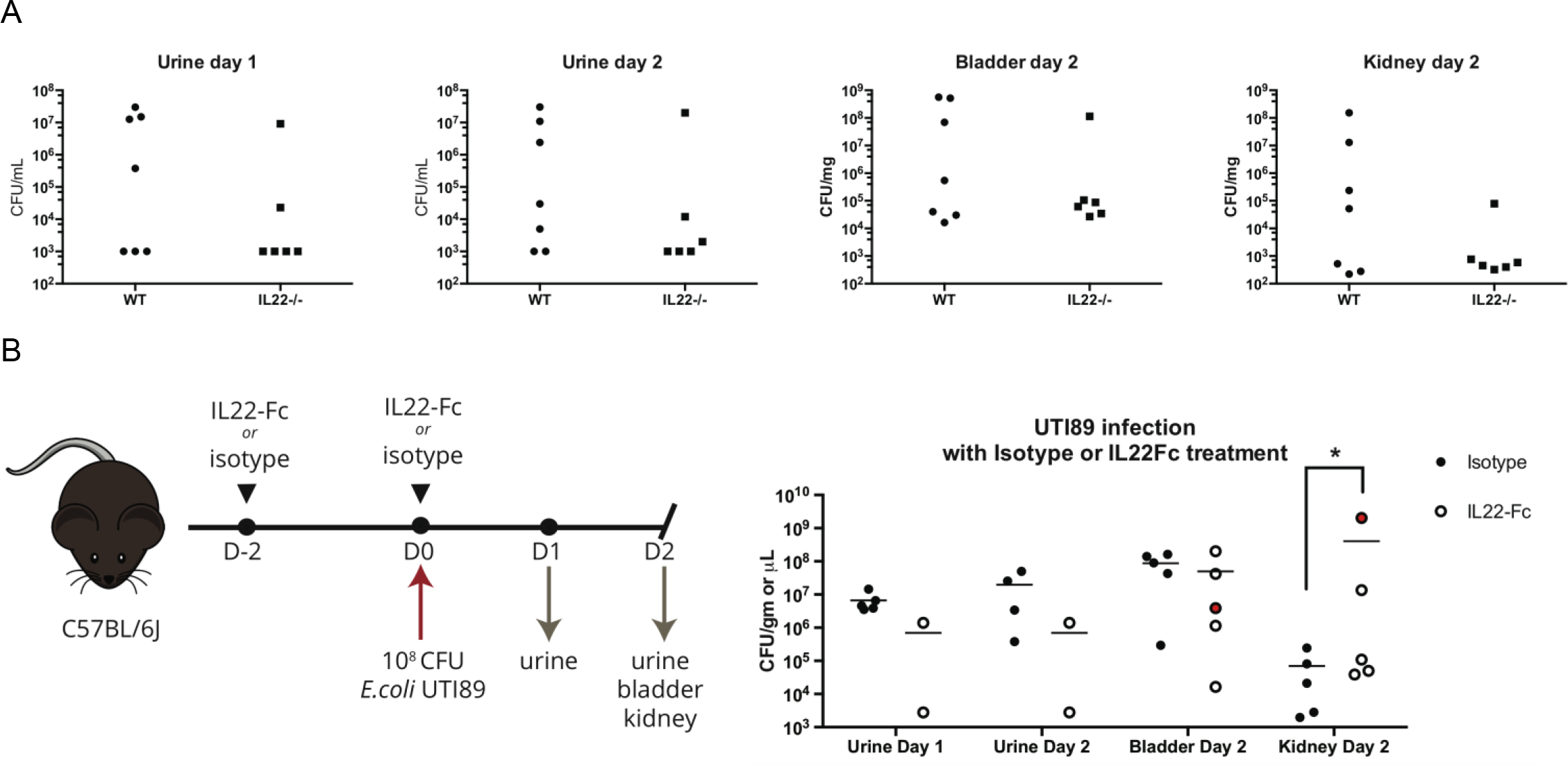
The effect of IL-22 on bacterial urinary tract infection. (A) *IL22*-/-and WT littermate female mice were transurethrally infected with 1×10^8^ CFU of *E.coli* UTI89. Shed urine was collected on day 1 and day 2, and on day 2 animals were euthanized for kidney and bladder CFU quantification. (B) 100μg IL-22-Fc (IL-22 cytokine conjugated to Fc to increase *in vivo* half life) or Fc isotype control (both courtesy of Genentech) were given to C57BL/6J female mice by IP injection at day −2 and immediately after transurethral infection with 1×10^8^ CFU of *E.coli* UTI89. Shed urine was collected on day 1 and day 2, and on day 2 animals were euthanized for kidney and bladder CFU quantification by plating on MacConkey agar. Several IL-22Fc treated animals shed little urine on days 1 and 2, and one IL22-Fc treated mouse (red dot) had to be euthanized on day 1 due to pyelonephritis-related morbidity.

## Discussion

The mechanisms by which the inflammatory response influences epithelial biology in the bladder are only partially understood. Given IL-22’s demonstrated role in immune responses and chronic pathologies in other organs, we sought to investigate how this cytokine might influence outcomes of infection in the bladder. We have documented urothelial expression of IL22Rα1, a partner with ubiquitously expressed IL10R2 that forms the functional IL-22 receptor in epithelial cells. We then tested the role of IL-22 using the infection models of *Schistosoma haematobium*, a leading cause of infection-associated bladder fibrosis and cancer globally, and bacterial urinary tract infections with a clinically-relevant uropathogenic *E. coli* strain.

During chronic inflammation of the bladder due to *Schistosoma haematobium* eggs, we observed *IL22-/-* mice have significant changes in expression of genes related to urothelial development and transferase activity. Hedgehog and Wnt developmental pathways play important roles in regeneration of the urothelium after repeat injury^8^, and bladder cancer arises from *Shh*-expressing basal cells in the bladder^9^. The reduced activity of these pathways in *IL22-/-* mice suggests that they may be protected from induction of bladder cancer, though this remains to be directly tested. It will be interesting to evaluate whether inadequate negative regulation of IL-22, such as in mice lacking the IL-22 binding protein (IL-22BP)^6^, may promote cancer formation in this model of *Schistosoma haematobium* infection.

During bacterial infection, our experiments with *IL22-/-* mice and using exogenous IL-22 suggest IL-22 deficiency may in fact protect from UTI89 infection. Given the potential influence of IL-22 on urothelial regeneration we observe during *S. haematobium* egg-induced inflammation, alterations of urothelial shedding and regeneration may underlie this difference. Shedding of the urothelium is an important innate defense against FimH expressing UTI89; FimH is part of the type 1 pilus that enables bacterial attachment to uroplakins present on the surface of mature urothelial cells^10^. Presumably, IL-22 may maintain uroplakin-expressing urothelial cells that serve as targets for colonization by UTI89, though this remains to be tested. Further, the effect of IL22-Fc on UTI89 infection may be indirect, given the influence of IL-22 on production of systemic acute phase response proteins^11^.

STAT3 signaling has been implicated downstream of IL-22. However, we have not observed global induction of phospho-STAT3 or other canonical signaling pathways in urothelial cells after *ex vivo* treatment of bladder tissue with IL-22 (data not shown), indicating further work is needed to understand the direct effects of IL-22 on urothelial functioning.

This report is the first examination of the potential roles of IL-22 in the bladder using two important bacterial infection models. Further experiments will identify the mechanisms by which IL-22 influences tissue homeostasis and defense during urinary tract infection. This will provide valuable insight into how the bladder mounts productive defense and repair responses and how these processes in chronic settings may lead to pathology. Moreover, our results suggest that IL-22 may be a therapeutic target for urinary tract infection.

## Methods

### Mice

We used 8-12 week old C57BL/6J and BALB/c female mice from Jackson Laboratories in accordance with protocols approved by the Institutional Animal Care and Use Committee (IACUC) at Stanford University School of Medicine. *IL22-/-* male mice on a B6 background were provided by Wenjun Ouyang at Genentech (South San Francisco, CA) and bred to C57BL/6J female mice in animal facilities at Stanford University. IL22+/-F1 progeny were interbred to generate F2 IL22-/-and WT littermates for exposure to *S. haematobium* and *E. coli* infection.

### Bladder wall injection

Bladder wall injections were performed as previously described.^12^ Briefly, *Schistosoma haematobium* eggs were isolated from the liver and intestinal tissues of a patently infected Syrian golden hamster. The bladder was exposed surgically, and 3000 eggs or vehicle alone were injected in a 50μL volume into the bladder wall.

### Urinary tract infection

Female mice were infected by transurethral catheterization^13^ with 1 × 10^8^ CFU of static overnight cultures of uropathogenic *Escherichia coli* UTI89. CFUs were determined by plating dilutions on MacConkey agar. Where indicated, mice were treated with 100μg IL22-Fc or isotype control (provided by Wenjun Ouyang at Genentech) by intraperitoneal injection.

### RNA isolation, qPCR, and microarray

Bladder tissue was homogenized by bead-beating with zirconium oxide beads in TRI Reagent (Ambion), and the RNA was precipitated with isopropanol and ethanol washes. RNA was further purified using the RNeasy MinElute Cleanup Kit (Qiagen). For qPCR, 1μg RNA was reverse transcribed using oligo-dT primers and the Superscript III RT kit (Invitrogen). qPCR for each gene was performed in triplicate using SYBR Select (Invitrogen) on a Stratagene MX3005p instrument. Primer sequences were obtained from PrimerBank: http://pga.mgh.harvard.edu/primerbank/ Relative expression calculated by the ΔΔC_t_ method with *Hprt or Gapdh* as the reference gene, as indicated. The Stanford Functional Genomics Facility (SFGF) processed RNA for hybridization to the Agilent Mouse V1 array, and data was analyzed using GeneSpring 12.6.1 software (Agilent).

### Immunofluorescence microscopy

Naïve C57BL/6J female mouse bladders were snap frozen in OCT. 5μm sections were fixed briefly in 2% PFA before blocking with 5% goat serum block. Sections were stained in 0.3% Triton X100 in PBS with rat anti-mouse IL-22Rα1 (RnD Systems), rabbit anti-cytokeratin 5 (Abcam), or isotype controls and goat secondary antibodies (Invitrogen). Slides were mounted in ProLong Gold with DAPI (Invitrogen) and imaged on a Nikon epifluorescent microscope.

### Histology

Mouse bladders were fixed for 24 hours in 10% buffered formalin, then dehydrated and embedded in paraffin. Five-micron sections were made stained with haematoxylin & eosin (H&E) for morphological assessment by a pathologist who evaluated the sections blindly.

## Acknowledgements

We gratefully thank Wenjun Ouyang for providing critical reagents.

